# Ensemble learning for robust knee cartilage segmentation: data from the osteoarthritis initiative

**DOI:** 10.1101/2020.09.01.267872

**Authors:** Edward J Peake, Raphael Chevasson, Stefan Pszczolkowski, Dorothee P Auer, Christoph Arthofer

## Abstract

**Purpose:** To evaluate the performance of an ensemble learning approach for fully automated cartilage segmentation on knee magnetic resonance images of patients with osteoarthritis.

**Materials and Methods:** This retrospective study of 88 participants with knee osteoarthritis involved the study of three-dimensional (3D) double echo steady state (DESS) MR imaging volumes with manual segmentations for 6 different compartments of cartilage (Data available from the Osteoarthritis Initiative). We propose ensemble learning to boost the sensitivity of our deep learning method by combining predictions from two models, a U-Net for the segmentation of two labels (cartilage vs background) and a multi-label U-Net for specific cartilage compartments. Segmentation accuracy is evaluated using Dice coefficient, while volumetric measures and Bland Altman plots provide complimentary information when assessing segmentation results.

**Results:** Our model showed excellent accuracy for all 6 cartilage locations: femoral 0.88, medial tibial 0.84, lateral tibial 0.88, patellar 0.85, medial meniscal 0.85 and lateral meniscal 0.90. The average volume correlation was 0.988, overestimating volume by 9% ± 14% over all compartments. Simple post processing creates a single 3D connected component per compartment resulting in higher anatomical face validity.

**Conclusion:** Our model produces automated segmentation with high Dice coefficients when compared to expert manual annotations and leads to the recovery of missing labels in the manual annotations, while also creating smoother, more realistic boundaries avoiding slice discontinuity artifacts present in the manual annotations.

**Key Results:**

- Combining a 2-label U-Net (cartilage vs background) with a multi-class U-Net for segmentation of cartilage compartment boosts the accuracy of our deep learning model leading to the recovery of missing annotations in the manual dataset.
- Automatically generated segmentations have high Dice coefficients (0.85-0.90) and reduce inter-slice discontinuity artefact caused by slice wise delineation.
- Model refinement yields more anatomically plausible segmentations where each cartilage label is composed of only a single 3D region of interest.

## 1. Introduction

Osteoarthritis (OA) is the most common joint disorder; the highest occurrence of OA is the knee joint with an estimated prevalence of 13% in the adult population in Europe (1,2). It is associated with an increasing socioeconomic impact owing to the aging and increasingly obese population (3). Manual segmentation of cartilage tissue from MRI has shown sufficient sensitivity to detect changes, including early progression (4,5). Also, greater rates of cartilage loss are associated with frequent knee pain, irrespective of adjustment or stratification for radiographic disease (6). While clinical OA is a late stage condition for which disease modifying opportunities are limited, early identification may offer a window of time for potentially more effective treatment (7). However, manual segmentation of cartilage is time-consuming and susceptible to inter-rater variability (8) while traditional automated registration and shape-based classification (9–13) have weaker performance metrics when compared to deep learning approaches (14–18).

Fully Convolutional Neural Networks (CNNs) provide an efficient and precise alternative for the segmentation of many anatomical structures in medical images. (19–21). In particular U-Nets, which comprise a contraction path (encoder) to capture context followed by an expansion path (decoder) to enable localisation, and shortcuts between selected layers, have shown improved segmentation accuracy (22).

Two dimensional and more resource-expensive fully volumetric 3D U-Nets have been proposed for multi-label segmentation of knee cartilage (15–17). U-Net-derived architectures have shown promising results when trained and evaluated on small datasets with as few as 20 subjects without augmentation for the segmentation of knee joint structures (9). Further refinements have been made using conditional random fields (22), statistical shape modelling (SSM), and applied to meniscal cartilage (23) femoral cartilage (11) and tibial cartilage (11,12). 2D U-Nets may struggle to differentiate medial from lateral cartilage. While SSM methods referenced may overcome this using 3D U-Nets and bespoke registration templates they are more computationally expensive.

In this study, we propose ensemble learning to increase the accuracy of our deep learning method while keeping computational resources limited by training a 2D U-Net with only two labels (cartilage vs background) to offset the high inter-rater variability in manual annotations (13,24), and a multi-label U-Net to identify cartilage compartment. Using simple post-processing, we create anatomically valid segmentations that contain only a single region of interest per cartilage label. Our ensemble learning approach performed as well as or better than existing methods that provide labels for all cartilage compartments.

## 2. Materials and Methods

### 2.1 Study Dataset

We utilized data from the Osteoarthritis Initiative (OAI), with cartilage annotations produced by iMorphics which are available online (25). The data for this retrospective analysis consisted of 3D-DESS MRI scans from 88 subjects with osteoarthritis; imaging was acquired at two timepoints doubling the number of images available for training. We included images from all subjects with manual segmentations for 6 cartilage compartments. The manual segmentations were performed by a trained musculoskeletal radiologist and reviewed by a technical specialist at iMorphics (26). The segmentations passed the iMorphics cartilage segmentation training protocol, requiring an intra-observer coefficient of variation lower than 3% on paired test images (26). The subjects consisted of 45 male and 43 female participants of age 61.2 ± 10.0 years and BMI of 31.3 ± 4.6kg/m^2^ (mean ± standard deviation). Based on the Kellgren-Lawrence (KL) scores for classification of knee OA (27), baseline scores were: KL-1 2%, KL-2 34%, KL-3 59% and KL-4 5%. MRI volumes had a 140mm field of view; 0.7mm slice thickness and matrix of 384×384 with 160 slices. Manual annotations were provided for the patellar, femoral, lateral tibial, medial tibial, lateral meniscal and medial meniscal cartilage.

### 2.2 Data Preparation

We applied N4 Bias field correction to remove low frequency intensity non-uniformity in the MRI data (28). Each MRI volume was intensity normalized by subtracting the mean voxel intensity and dividing by the standard deviation. As the dataset consisted of baseline and 12-month MRI we chose to randomly split the dataset by subject to avoid the same person appearing in the training and test sets. We used a 70/20/10% split for training, validation, and testing. To train our 2-label U-Net (2l model) for cartilage segmentation we reduced the number of labels to two, cartilage and background. For the 7-label U-Net (7l model) we used the fully annotated dataset as provided by iMorphics.

### 2.3 Model Architecture and training

Using the U-Net architecture shown in Figure 1, we trained a 2-label model for segmentation of cartilage from background and a 7-label model for segmentation of each cartilage compartment. Convolutional layer inputs were zero padded to preserve the height and width of the outputs from these layers. We trained our network using the Adam optimizer (29) with a learning rate of 1×10^−4^; a weighted categorical cross-entropy loss function was used to account for imbalances in the number of training examples for each cartilage label. We trained the U-Net for 300 epochs on the training dataset. At each epoch, the Dice coefficients were averaged over all labels for the entire validation dataset to give a multi-label Dice score. The model weights from the epoch with the highest multi-label Dice coefficient was selected for our combined model.

**Figure 1:**
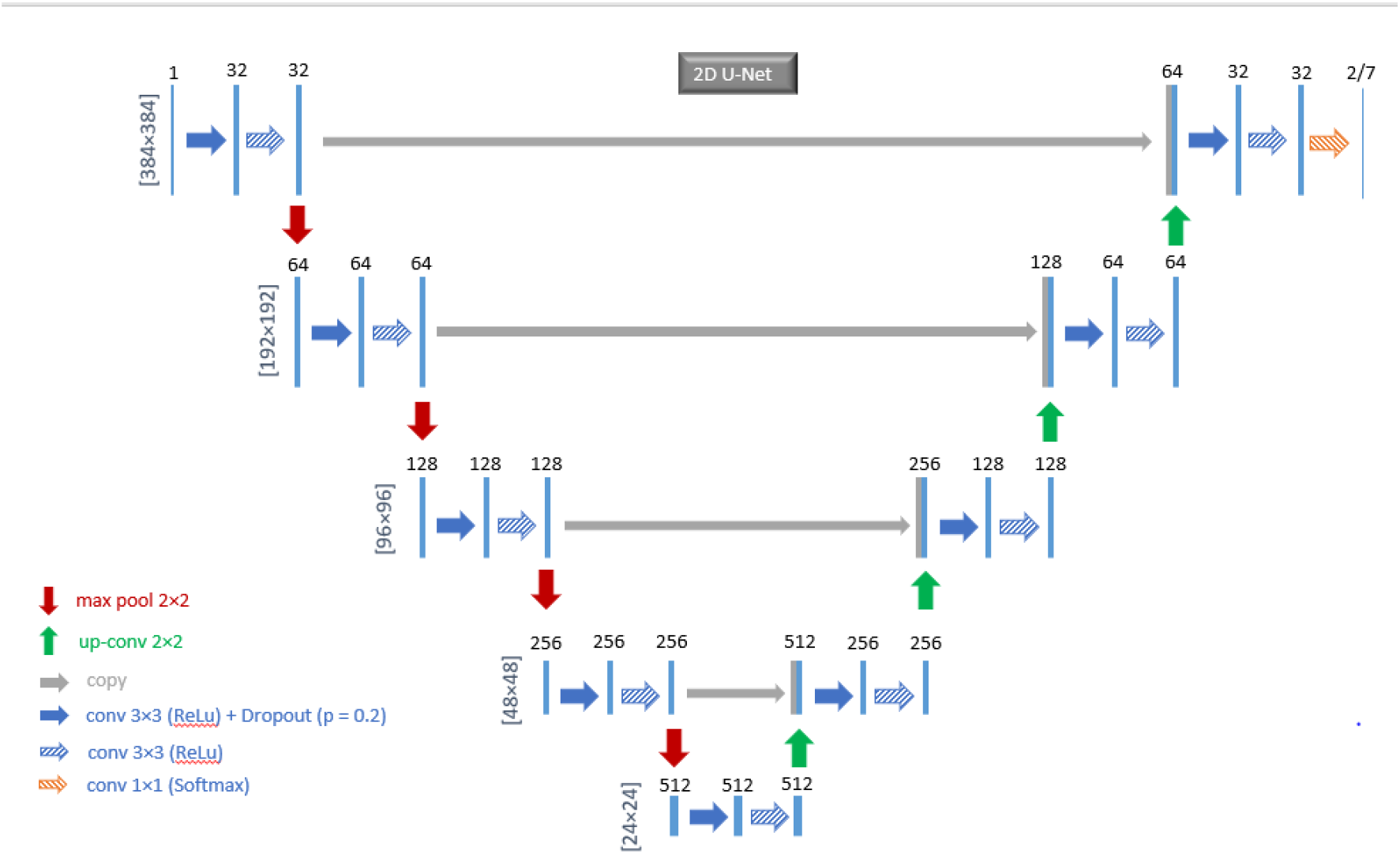
U-net architecture for cartilage segmentation. Each blue box corresponds to a multi-channel feature map. The number of channels is denoted on top of the box. The x-y-size is provided at the left edge of each depth layer. Grey boxes represent copied feature maps. The arrows denote the different operations. In the expansion path the first 3 ×3 Convolution operator has a stride of 2 to reduce the size of the feature map. In the last output operation (softmax layer) the number of channels is changed to train either a 2-label cartilage model or a 7-lable cartilage model.

We combined our models based on pixel voting; the 2-label model was used to determine if a voxel contained cartilage, and if so the 7-label model was used to determine the cartilage compartment (ignoring the background class). All code was implemented in Python 3.7.3 utilizing Tensorflow 1.13.1 (https://www.tensorflow.org/) and Keras 2.2.4 (https://keras.io/) packages. All training and evaluation were carried out on a dedicated server hosting a 64-bit CentOS 7 operating system. Computing hardware included 80 Intel Xeon E5-2698 v4 CPUs @ 2.20GHz, 512GB DDR4RAM, and two Nvidia Tesla K10 graphic cards.

### 2.4 Post-processing

To refine the outputs of our combined network (7l2l model) we applied rule-based post processing to the segmentation maps provided by our model (Figure 2). For each cartilage compartment, we relabelled the connected components based on the labels of neighbouring voxels where the largest connected component keeps its original labels, assigned by the U-Net. We used 3D connected components with a connectivity type of 26 such that voxels were connected if their faces, edges, or corners touch. The remaining connected components were relabelled based on the most frequent label of neighbouring voxels.

**Figure 2:**
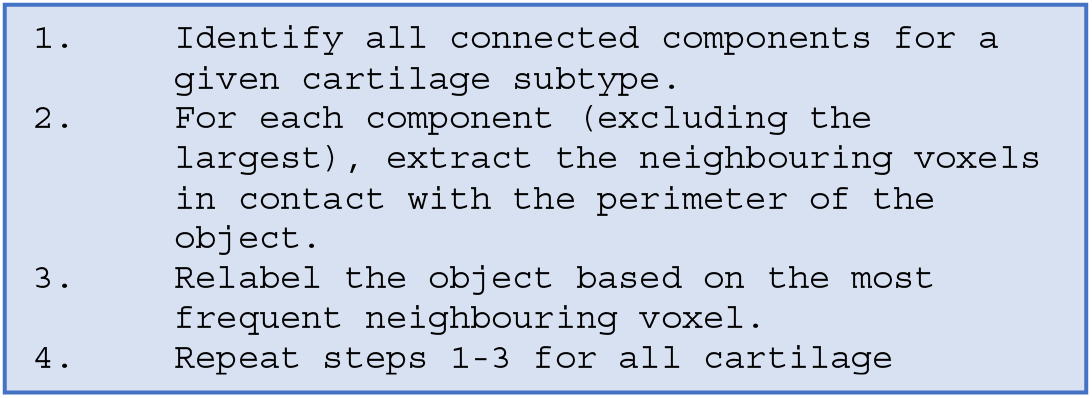
Refinement of the combined model, mislabelled cartilage is either relabelled or removed. This post processing reduces the number of connected components to a single connected component per cartilage type.

### 2.5 Model Evaluation

Evaluation was performed based on a 3-fold cross validation approach in which 10% of the data was held out for testing. We took a similar approach to previous studies which used a varying number of folds ranging from 2 to 5 folds (16,17,23). For each fold we used the remaining 70% for training the model and 20% for internal validation at each epoch to prevent overfitting. The resulting segmentations from the test sets were pooled when evaluating our segmentation model. The accuracy of tissue segmentation was calculated using the Dice similarity coefficient for each cartilage compartment, and was defined as DSC = 2|S∩R|/(|S|+|R|) where S denotes the automatically segmented voxels and R denotes the reference manually annotated voxels. Dice coefficients have a range of between 0 and 1 for each cartilage compartment with 1 representing a perfect overlap and 0 representing no overlap of the segmentations. Volumetric measures (scatter and Bland-Altman) were used to compare our corrected 7l2l model and manual segmentations.

## 3. Results

For each fold the overall training time was approximately 48 hours given the computing hardware in the current study. The segmentation of all 6 regions of interest took 13.6s in total for each image volume with a mean computing time of 3.6s for the application of the combined U-Net and 10s for the post-processing.

Figure 3 shows the Dice coefficients for each individual segmented cartilage structure. The Dice coefficient for each label was evaluated over the test sets obtained from the 3-fold cross validation; this was to avoid small or large cartilage regions from a given subject from biasing the Dice score, for a given cartilage compartment. Our combined 7l2l model achieved higher Dice scores across all compartments compared to our initial 7l model. While the improvements in Dice from post processing were small (corrected 7l2l model) the number of connected components was reduced from 21 per cartilage compartment to just one connected component per cartilage compartment. Visual assessment showed that the refinement yields more anatomically plausible segmentations, which may improve the quality of radiomics features from these segmentations.

**Figure 3:**
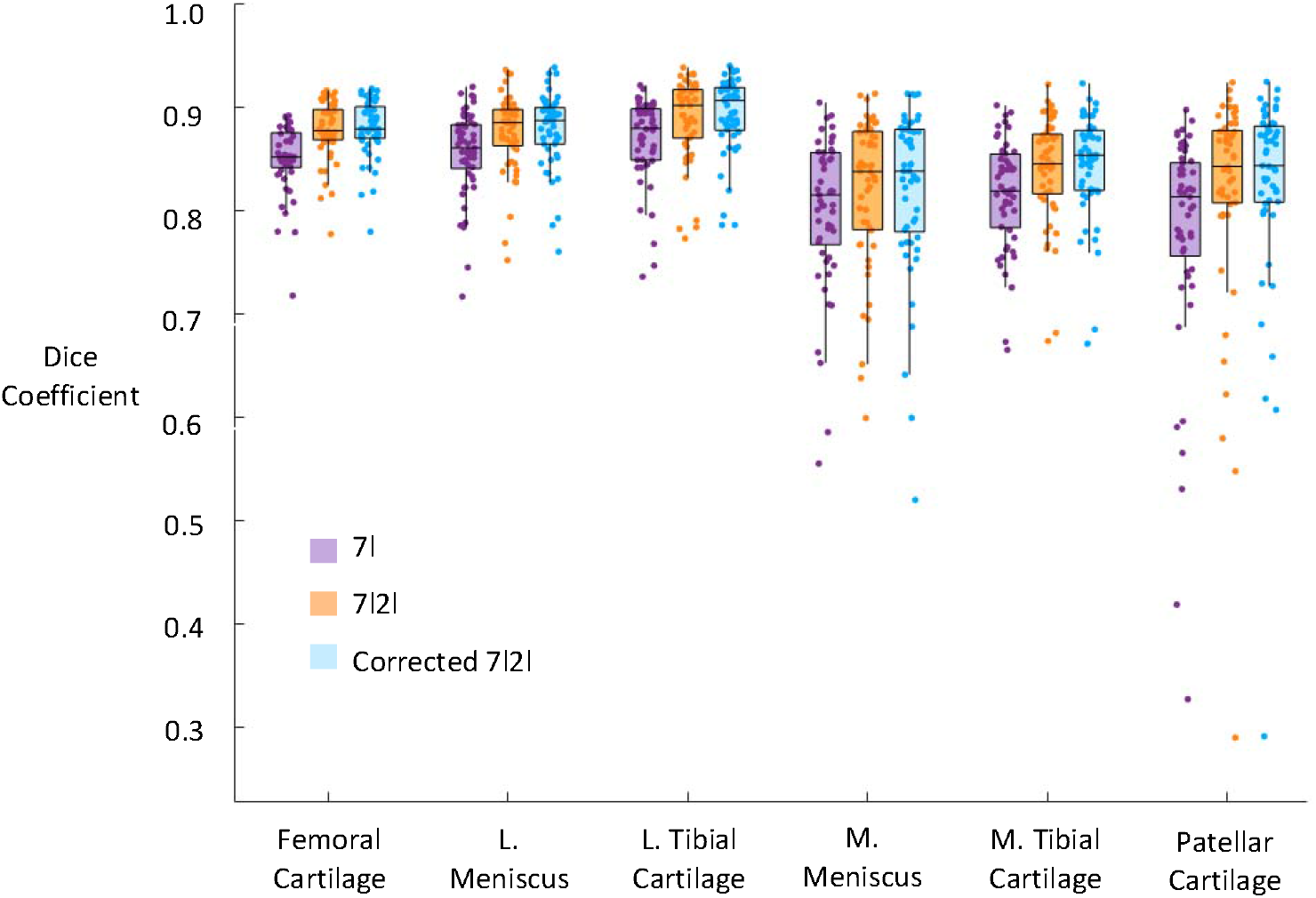
Box and jitter plot of the Dice coefficient values for each segmented cartilage type in the pooled test sets. Values are shown for the 7 label U-net (7l), combined model (7l2l), and corrected 7l2l model. Dice scores have a higher value and have a lower standard deviation for femoral cartilage, L. meniscal and L. tibial cartilage compared to patellar, M. meniscal and M. tibial cartilage.

The Dice coefficient was used to compare our results to other automated methods for segmentation using the iMorphics dataset (Table 1). Our approach reached state-of-the-art segmentation accuracy compared to other deep learning based methods (11,12,15–18,23) and outperformed competing methods that also provided automated segmentations for all cartilage labels. Dice coefficients (mean and 95% confidence intervals) for our method were: femoral 0.88 ± 0.03, medial tibial 0.84 ± 0.08, lateral tibial 0.88 ± 0.04, patellar 0.85 ± 0.10, medial meniscal 0.85 ± 0.05 and lateral meniscal 0.90 ± 0.04.

**Table 1:**
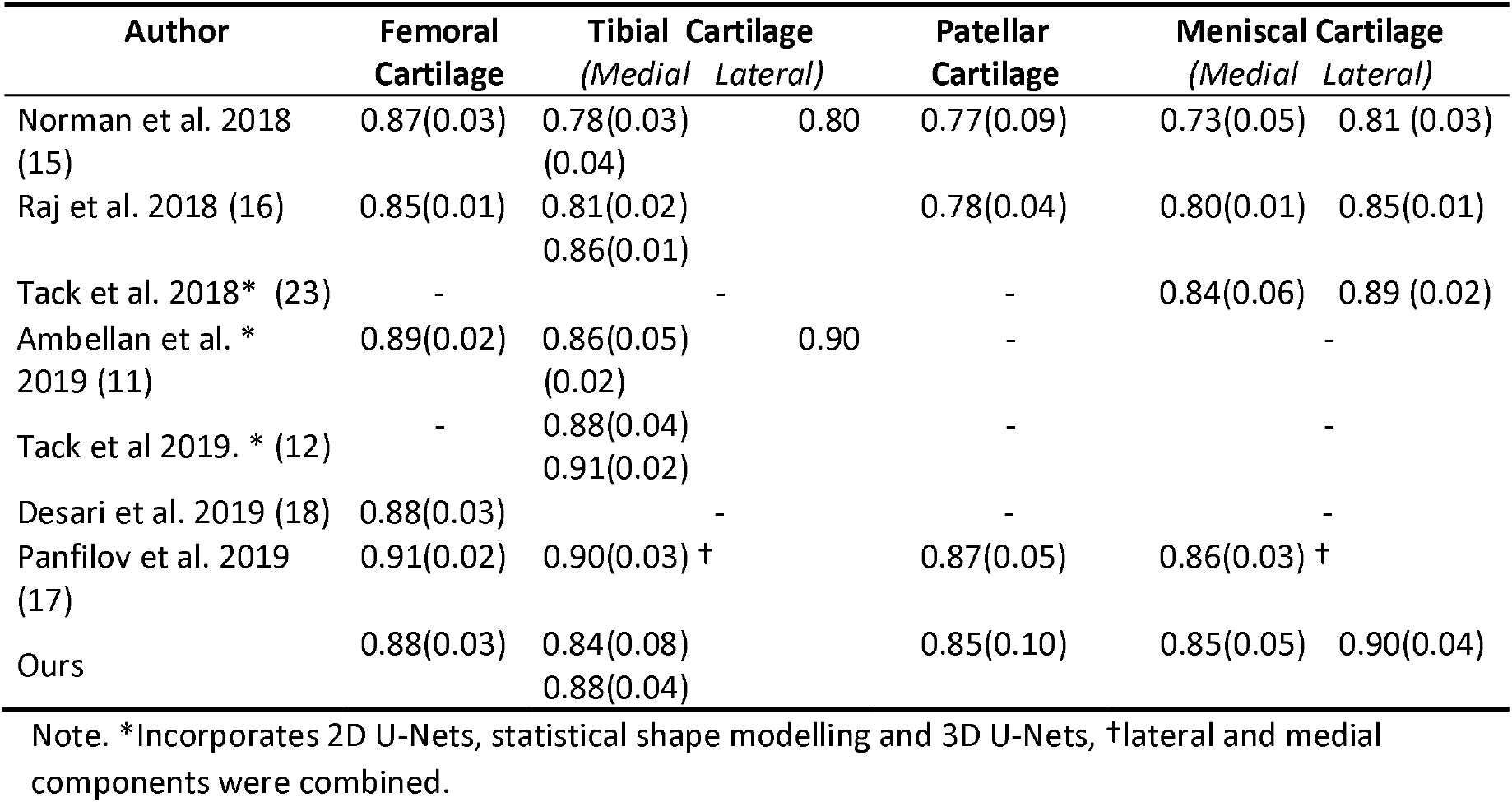
Dice coefficients for automatic segmentation methods using the iMorphics dataset. Data are means with 95% intervals in parentheses.

Figure 4 shows sagittal examples of tissue segmentation performed on the 3D DESS images in the test sets. The results from the 7-label U-Net demonstrate good agreement with the overall contours of the manual segmentation. In the example, the annotation for the lateral meniscus is missing in the manual annotations. The intermediary (uncorrected) 7l2l model recovers some of this information but does not correctly label all the voxels; the proper label is recovered by our corrected 7c2c model.

**Figure 4.**
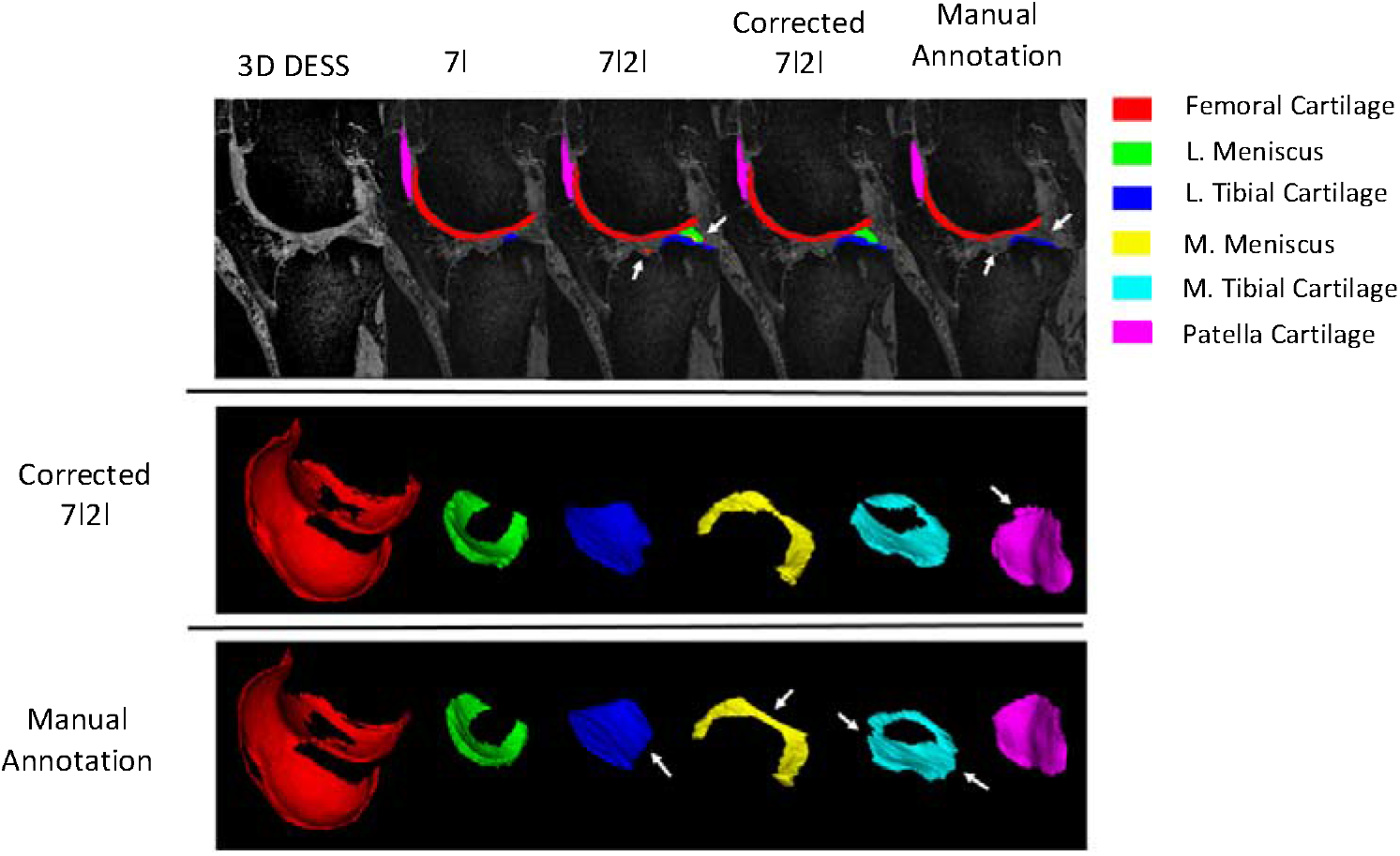
Tissue segmentations performed on a 3D Double Echo Stead State (DESS) MRI of a subject with full thickness cartilage loss in the medial compartment. Sagittal slices show the different models, while the 3D renderings are generated for the corrected 7l2l model and manual segmentations; these are visually very similar and demonstrate similar patterns of cartilage damage. Note the recovery of some unlabeled lateral meniscal cartilage in our corrected 7l2l model, and smoother border profiles in the 3D renderings.

Also shown in Figure 4 are the 3D rendered volumes for the corrected 7l2l model and manual segmentation. Noticeably, in the presence of advanced tissue degeneration, there is less agreement between the automatically segmented structures and the manual annotations. This is partly due to the increased subjectivity of segmentation in these regions as adjacent structures have comparable signal contrasts. In addition, cropping and irregular boundaries are visible in the manual annotations, while the automated segmentations have smoother boundaries. However, automated segmentations occasionally spill over into adjacent structures, as seen in the patellar.

Figure 5 shows scatter and Bland-Altman plots for our corrected 7l2l model when applied to the pooled test sets. The volume of test data sets for manual and automatic segmentations show a strong linear relationship across all cartilage compartments. For the pooled test sets, the average volume correlation across cartilage compartments was 0.988, with an average absolute mean difference across compartments of 0.43cm^3^. The regression on the scatterplot shows a significant trend towards larger predicted volumes (gradient 1.034, 95% confidence interval 1.003 – 1.067). On the Bland Altman plot, the root mean squared error (RMSE) appears stable across the range of volumes (−9.3%). While the standard deviation is normally distributed (Kolmogorov–Smirnov test, p < 0.05) the percentage difference between actual and predicted volume tends to decrease for larger volumes.

**Figure 5.**
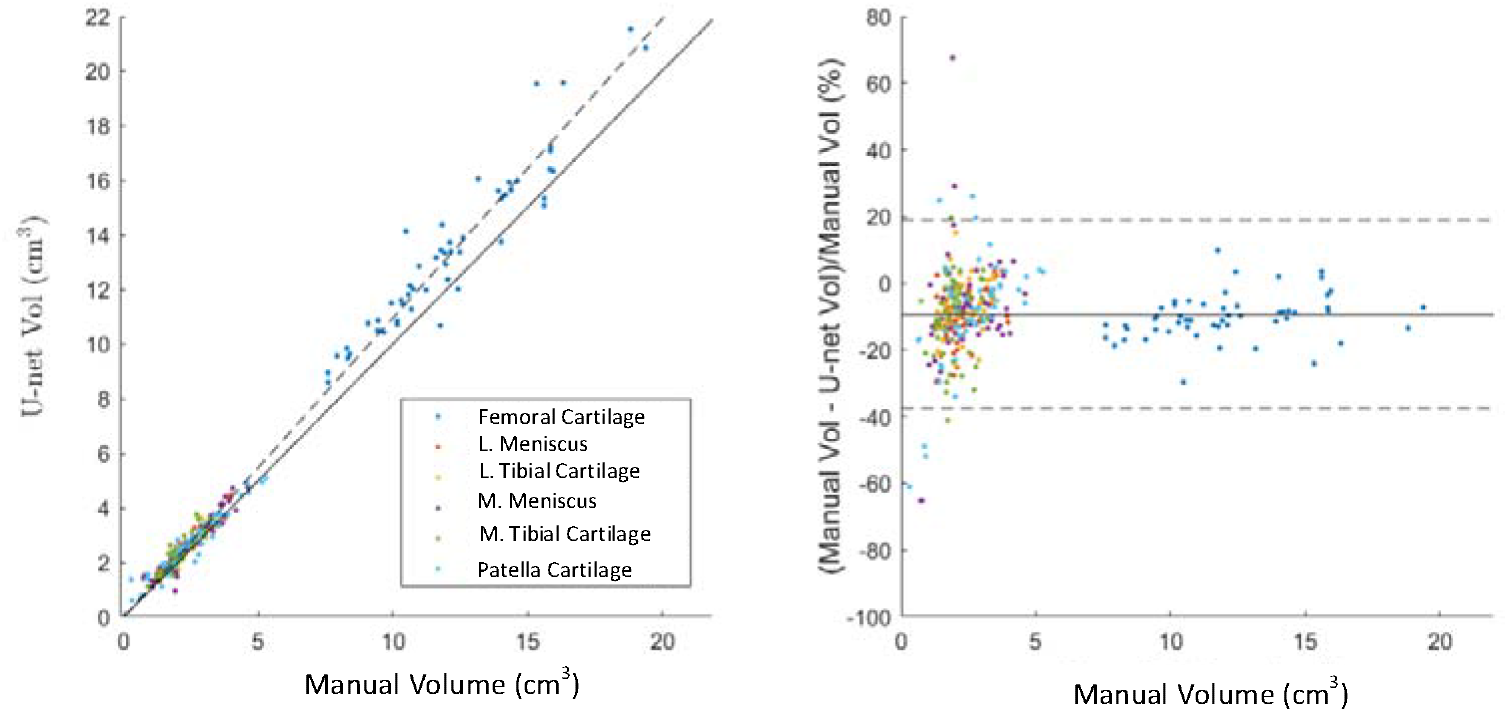
Scatterplots and Bland-Altman plot shows the comparison of volumetric measurements produced from our corrected 7l2l model compared to manual annotations, results are broken down by cartilage type. (Note that the linear regression, mean difference, and standard errors were calculated using the entire test data, not between compartments.)

## 4. Discussion

In this study we aimed to increase the accuracy of automated knee cartilage segmentation by combining predictions from a 2 label (cartilage vs background) U-Net with a 7-label U-Net to determine the type of cartilage. Our results achieved the highest Dice scores of any competing models that provide segmentations for all 6-cartilage compartments. Authors used different splits of training, validation, and testing so results should be compared carefully. We demonstrate a gain in Dice across all cartilage labels when comparing our combined model to our 7-label model. In addition, our post-processing is very simple and has a similar effect to SSM in removing small isolated regions in the segmentation mask and assigning the correct labels to different cartilage compartments.

Studies that have used SSM to provide segmentations for specific cartilage targets have also produced high Dice scores (11,12,23). However, this multistep process requires 2D and 3D U-Nets in addition to prior shape information for each target; also this method has been shown to fail to adequately segment cases which exhibit full thickness cartilage denudation (23). Furthermore, SSM methods are more computationally expensive; based on estimates using a single computational node segmentation of a single knee joint (cartilage and bone) took around 10mins (11). Despite differences in hardware U-nets are computationally more efficient, our combined model takes 13.6s to segment a single MRI volume, making it more feasible to segment entire databases such as the OAI which contains 50000 or more MRIs.

We observe that Dice coefficients of medial cartilage is lower than that of their lateral counterparts; a finding reproduced in other studies (11,12,15,16). The prevalence of radiographic medial compartment OA may be 5–10 times higher than in the lateral compartment (30). We hence suggest that lower Dice coefficients may result from greater uncertainty in the manual segmentations in medial regions due to advanced cartilage degeneration. Notably, for manual segmentations where the volume is less than 50% of the mean for a given label (n = 14) the average Dice coefficient is 0.68, compared to remainder of the data (n = 304) with a DICE coefficient of 0.87.

The Dice coefficient for inter-rater agreement was 0.79 in a similar dataset (24), while we achieved a Dice coefficient ≥ 0.85 across all cartilage compartments; suggesting automated segmentation is at least as good as manually generated annotations. When comparing our corrected 2l7l U-net results to those presented by Normal et al (15) the average volume correlation is higher (0.988 compared to 0.938), and the average absolute mean difference across compartments is lower (compared to 0.510cm^3^).

Our U-net architecture is 2D rather than 3D and therefore does not account for spatial information in the through-slice direction. This reduces the constrain to match the shape of the manual annotations in this direction which exhibit inter-slice discontinuity artefacts caused by slice wise delineation, resulting in the recovery of missing labels in the manual annotations, and smoother, more realistic boundaries around automatically generated segmentations. When compared to the 2D U-net by Panfilov et al (17), our Dice coefficients are slightly lower, however our model is able to differentiate medial from lateral components which may be of particular importance given the role of laterality in knee osteoarthritis.

U-nets tend to overestimate total volume segmentation in the OAI dataset, in the study by Dam et al. this was 14% for medial tibial and femoral compartments (31), and 12% across all compartments measured by Normal et al (8). In our study, or corrected 2l7l U-net overestimated the volume by 9% over all compartments. This result is similar to inter-observer reproducibility measures by Kristina et al who calculated a RMSE of 9.9% for medial meniscal cartilage using DESS MRI volumes from the OAI (32).

In this study we do not have quantifiable measures as to the causes of the differences between manual annotations and automatic segmentations. This limitation could be addressed in further work using higher resolution MRI, utilizing either higher field strengths or post-mortem imaging. Our model is limited by its lack of generalizability to clinical imaging protocols as training data consists entirely of DESS MRI. Improvements could include augmenting training data from clinically acquired scans or using generative adversarial networks to create images with synthetic DESS contrast.

In this paper, we propose an ensemble learning approach based on a 2-label 2D U-Net to differentiate cartilage from background, followed by a multi-label U-Net to distinguish between different cartilage types. Our combined model offers rapid and accurate cartilage segmentation with favourable performance compared to previous multi-compartment segmentation algorithms that were tested using the OAI/iMorphics dataset. Visually, the presented automated segmentations appear smoother with anatomically more realistic boundaries compared to manual annotations.

## Acknowledgment

Data and/or research tools used in the preparation of this manuscript were obtained and analyzed from the controlled access datasets distributed from the Osteoarthritis Initiative (OAI), a data repository housed within the NIMH Data Archive (NDA). OAI is a collaborative informatics system created by the National Institute of Mental Health and the National Institute of Arthritis, Musculoskeletal and Skin Diseases (NIAMS) to provide a worldwide resource to quicken the pace of biomarker identification, scientific investigation and OA drug development.

## References

1. Pereira D, Peleteiro B, Araújo J, Branco J, Santos RA, Ramos E. The effect of osteoarthritis definition on prevalence and incidence estimates: A systematic review [Internet]. Vol. 19, Osteoarthritis and Cartilage. 2011 [cited 2020 Mar 23]. p. 1270–85. Available from: http://www.ncbi.nlm.nih.gov/pubmed/21907813

2. Prieto-Alhambra D, Judge A, Javaid MK, Cooper C, Diez-Perez A, Arden NK. Incidence and risk factors for clinically diagnosed knee, hip and hand osteoarthritis: Influences of age, gender and osteoarthritis affecting other joints. Ann Rheum Dis [Internet]. 2014 Sep [cited 2020 Mar 23];73(9):1659–64. Available from: http://www.ncbi.nlm.nih.gov/pubmed/23744977

3. Hunter DJ, Bierma-Zeinstra S. Osteoarthritis. Vol. 393, The Lancet. Lancet Publishing Group; 2019. p. 1745–59.

4. Eckstein F, Cicuttini F, Raynauld JP, Waterton JC, Peterfy C. Magnetic resonance imaging (MRI) of articular cartilage in knee osteoarthritis (OA): morphological assessment. Osteoarthr Cartil. 2006;14(SUPPL. 1):46–75.

5. Wang Y, Wluka AE, Cicuttini FM, Jones G, Ding C. Use magnetic resonance imaging to assess articular cartilage. Vol. 4, Therapeutic Advances in Musculoskeletal Disease. SAGE Publications; 2012. p. 77–97.

6. Eckstein F, Cotofana S, Wirth W, Nevitt M, John MR, Dreher D, et al. Greater rates of cartilage loss in painful knees than in pain-free knees after adjustment for radiographic disease stage: Data from the osteoarthritis initiative. Arthritis Rheum [Internet]. 2011 Aug [cited 2020 Apr 21];63(8):2257–67. Available from: http://www.ncbi.nlm.nih.gov/pubmed/21520009

7. Chu CR, Williams AA, Coyle CH, Bowers ME. Early diagnosis to enable early treatment of pre-osteoarthritis. Vol. 14, Arthritis Research and Therapy. BioMed Central; 2012. p. 212.

8. Gatti AA. NEURALSEG: state-of-the-art cartilage segmentation using deep learning – analyses of data from the osteoarthritis initiative. Osteoarthr Cartil [Internet]. 2018 Apr 1 [cited 2019 Jul 29];26:S47–8. Available from: https://www.sciencedirect.com/science/article/pii/S1063458418302103

9. Zhou Z, Zhao G, Kijowski R, Liu F. Deep convolutional neural network for segmentation of knee joint anatomy. Magn Reson Med [Internet]. 2018 Dec [cited 2019 Jul 29];80(6):2759–70. Available from: http://www.ncbi.nlm.nih.gov/pubmed/29774599

10. Tack A, Mukhopadhyay A, Zachow S. Knee menisci segmentation using convolutional neural networks: data from the Osteoarthritis Initiative. Osteoarthr Cartil [Internet]. 2018 May 1 [cited 2019 Jul 29];26(5):680–8. Available from: https://www.sciencedirect.com/science/article/pii/S1063458418310811

11. Ambellan F, Tack A, Ehlke M, Zachow S. Automated segmentation of knee bone and cartilage combining statistical shape knowledge and convolutional neural networks: Data from the Osteoarthritis Initiative. Med Image Anal. 2019 Feb 1;52:109–18.

12. Tack A, Zachow S. Accurate Automated Volumetry of Cartilage of the Knee using Convolutional Neural Networks: Data from the Osteoarthritis Initiative. 2019;

13. Siorpaes K, Wenger A, Bloecker K, Wirth W, Hudelmaier M, Eckstein F. Interobserver reproducibility of quantitative meniscus analysis using coronal multiplanar DESS and IWTSE MR imaging. Magn Reson Med [Internet]. 2012 May [cited 2020 Mar 31];67(5):1419–26. Available from: http://www.ncbi.nlm.nih.gov/pubmed/22135245

14. Prasoon A, Petersen K, Igel C, Lauze F, Dam E, Nielsen M. Deep feature learning for knee cartilage segmentation using a triplanar convolutional neural network. In: Lecture Notes in Computer Science (including subseries Lecture Notes in Artificial Intelligence and Lecture Notes in Bioinformatics). 2013. p. 246–53.

15. Norman B, Pedoia V, Majumdar S. Use of 2D U-net convolutional neural networks for automated cartilage and meniscus segmentation of knee MR imaging data to determine relaxometry and morphometry. Radiology [Internet]. 2018 Jul 27 [cited 2019 Jul 29];288(1):177–85. Available from: http://pubs.rsna.org/doi/10.1148/radiol.2018172322

16. Raj A, Vishwanathan S, Ajani B, Krishnan K, Agarwal H. Automatic knee cartilage segmentation using fully volumetric convolutional neural networks for evaluation of osteoarthritis. In: Proceedings - International Symposium on Biomedical Imaging. IEEE Computer Society; 2018. p. 851–4.

17. Panfilov E, Tiulpin A, Klein S, Nieminen MT, Saarakkala S. Improving Robustness of Deep Learning Based Knee MRI Segmentation: Mixup and Adversarial Domain Adaptation. In: 2019 IEEE/CVF International Conference on Computer Vision Workshop (ICCVW) [Internet]. IEEE; 2019 [cited 2020 Mar 15]. p. 450–9. Available from: https://ieeexplore.ieee.org/document/9022164/

18. Desai AD, Gold GE, Hargreaves BA, Chaudhari AS. Technical Considerations for Semantic Segmentation in MRI using Convolutional Neural Networks. 2019 [cited 2020 Mar 15]; Available from: http://oai.epi-ucsf.org

19. Novikov AA, Lenis D, Major D, Hladůvka J, Wimmer M, Bühler K. Fully Convolutional Architectures for Multi-Class Segmentation in Chest Radiographs.

20. Wu S, Li H, Quang D, Guan Y. Three-Plane–assembled Deep Learning Segmentation of Gliomas. Radiol Artif Intell. 2020 Mar 1;2(2):e190011.

21. Çiçek Ö, Abdulkadir A, Lienkamp SS, Brox T, Ronneberger O. 3D U-net: Learning dense volumetric segmentation from sparse annotation. In: Lecture Notes in Computer Science (including subseries Lecture Notes in Artificial Intelligence and Lecture Notes in Bioinformatics). Springer Verlag; 2016. p. 424–32.

22. Ronneberger O, Fischer P, Brox T. U-Net: Convolutional Networks for Biomedical Image Segmentation [Internet]. [cited 2019 Jul 9]. Available from: http://lmb.informatik.uni-freiburg.de/

23. Tack A, Mukhopadhyay A, Zachow S. Knee menisci segmentation using convolutional neural networks: data from the Osteoarthritis Initiative. Osteoarthr Cartil [Internet]. 2018 May [cited 2019 Jul 29];26(5):680–8. Available from: https://linkinghub.elsevier.com/retrieve/pii/S1063458418310811

24. Heimann T, Morrison BJ, Styner MA, Niethammer M, Warfield SK. Segmentation of Knee Images: A Grand Challenge [Internet]. [cited 2020 Mar 16]. Available from: www.ski10.org

25. OAI:DATA FROM 3D CARTILAGE/MENISCUS SEGMENTATIONS OF KNEE MRI SCANS [Internet]. [cited 2020 Apr 2]. Available from: https://oai.epi-ucsf.org/datarelease/iMorphics.asp

26. Paproki A, Engstrom C, Chandra SS, Neubert A, Fripp J, Crozier S. Automated segmentation and analysis of normal and osteoarthritic knee menisci from magnetic resonance images--data from the Osteoarthritis Initiative. Osteoarthritis Cartilage. 2014 Sep 1;22(9):1259–70.

27. Kellgren JH, Lawrence JS. Radiological assessment of osteo-arthrosis. Ann Rheum Dis. 1957;16(4):494–502.

28. Tustison NJ, Avants BB, Cook PA, Yuanjie Zheng, Egan A, Yushkevich PA, et al. N4ITK: Improved N3 Bias Correction. IEEE Trans Med Imaging [Internet]. 2010 Jun [cited 2019 Jul 9];29(6):1310–20. Available from: http://ieeexplore.ieee.org/document/5445030/

29. Kingma DP, Ba J. Adam: A Method for Stochastic Optimization. 2014 Dec 22 [cited 2019 Aug 2]; Available from: http://arxiv.org/abs/1412.6980

30. Ahlbäck S. Osteoarthrosis of the knee. A radiographic investigation. Acta Radiol Diagn (Stockh). 1968;

31. Dam EB, Lillholm M, Marques J, Nielsen M. Automatic segmentation of high- and low-field knee MRIs using knee image quantification with data from the osteoarthritis initiative. J Med Imaging. 2015 Apr 20;2(2):024001.

32. Siorpaes K, Wenger A, Bloecker K, Wirth W, Hudelmaier M, Eckstein F. Interobserver reproducibility of quantitative meniscus analysis using coronal multiplanar DESS and IWTSE MR imaging. Magn Reson Med [Internet]. 2012 May [cited 2020 Aug 5];67(5):1419–26. Available from: pmc/articles/PMC3527078/?report=abstract

